# Monocular vision is intrinsically unstable: a side-effect of binocular homeostasis

**DOI:** 10.1101/2020.03.17.987362

**Authors:** Alexandre Reynaud, Kévin Blaize, Frédéric Chavane, Robert F. Hess

## Abstract

It is now accepted that short-term deprivation of one eye in adults results in not only a post-deprivation strengthening of the vision in the previously deprived eye but also a deterioration in the vision of the previously non-patched eye. Such monocular deprivation of 1-2 hours induces changes that last approximately 30-90 minutes. There is some support for this neuroplastic effect being the consequence of a change in the contrast gain within the binocular circuity. What is not known is when these changes in gain are initiated. One possibility is that they are initiated only once the patch is removed. The other possibility is that they are the result of a slow build up from the moment the patch is first applied.

In this study, we measure monocular contrast detection thresholds of the non-deprived eye over time during the deprivation of the other eye. We show that contrast threshold increases over time during the deprivation of the other eye. This observation suggest that the patching effect is mediated by a slow build up over the deprivation period: reducing the vision of the non-deprived eye and enhancing the vision of the deprived eye reflecting reciprocal changes in sensitivity. These results highlight a hitherto unknown feature of human vision, namely that monocular vision *per se* is intrinsically unstable which is a consequence of the reciprocal inhibitory circuits that homeostatically regulate binocular vision. This questions a whole corpus of studies of visual function that rely on the assumption that monocular vision is intrinsically stable.

## Introduction

It is now accepted that short-term deprivation of one eye in adults results in not only a post-deprivation strengthening of the vision in the previously deprived eye (Lunghi et al., 2011; Zhou et al., 2013) but also a deterioration in the vision of the previously non-patched eye (Zhou et al., 2013). Such monocular deprivation of 1-2 hours induces changes that last approximately 30-90 minutes. There is some support for this neuroplastic effect being the consequence of a change in the contrast gain within the binocular circuity (Lunghi et al., 2011;Zhou et al, 2013; but also see Binda et al., 2018). What is also not known is when these changes in gain are initiated. One possibility is that they are initiated only once the patch is removed, this would explain why the duration of deprivation has so little effect (Min et al., 2018). The other possibility is that they are the result of a slow build up from the moment the patch is first applied. If the latter is true, one consequence would be that vision under monocular conditions would be expected to be unstable. If true, this would undermine a great deal of what we presently know about human vision which has been obtained under monocular testing conditions where it has been assumed vision is stable, though never been critically tested. Here we show that short-term neuroplasticity builds up during the period of deprivation and as a consequence there is a slow deterioration of vision of the non-patched eye, resulting in monocular vision being inherently unstable over time.

In this study, we measure monocular contrast detection thresholds of the non-deprived eye over time during the deprivation of the other eye. We show that contrast threshold increases over time during the deprivation of the other eye. This observation suggest that the patching effect is mediated by a slow build up over the deprivation period: reducing the vision of the non-deprived eye and enhancing the vision of the deprived eye reflecting reciprocal changes in sensitivity.

## Material and Methods

### Apparatus

The experiment was programmed in Matlab (version R2016b) using the Psychtoolbox (Brainard, 1997; Pelli 1997; Kleiner et al., 2007) and a Bits# Stimulus Processor (Cambridge Research Systems Ltd., Kent, UK) in order to present stimuli with a 14-bit luminance resolution. Stimuli were displayed on a Dell P1130 monitor with a resolution of 1280 × 1024 pixels, a framerate of 85 Hz and a mean luminance of 82 cd/m^2^ in an artificially lit room. Subjects were asked to place their head on a chin-rest in order to ensure a constant viewing distance of 63 cm in each block.

### Participants

Fifteen subjects (6 males, 1 author, mean age 24.5 ± 5.0) with normal or corrected-to-normal vision and normal binocular vision (disparity threshold less than 100 arcmin, randot test) participated in the study. Informed consent was obtained from all subjects. This research has been approved by the Ethics Review Board of the McGill University Health Center and was performed in accordance with the ethical standards laid down in the Code of Ethics of the World Medical Association (Declaration of Helsinki). Informed consent was obtained from all subjects.

### Procedures

Two conditions were tested: binocular i.e. the control condition or monocular i.e. the deprived condition at two different spatial frequencies 1 c/d or 5 c/d for a total of 4 experimental sessions, tested on different days in a pseudo-randomized counterbalanced order. Threshold measurements were repeated at 0, 5, 10, 15, 30 and 60 minutes in the time course of the experiment. In the monocular condition, participants wore a translucent eye patch on one of their eyes during the whole one-hour course of the experiment.

Contrast thresholds were measured with an identification task using a staircase procedure (2 up - 1 down), with contrast chosen on a log-scale between 0.001 and 1. Initial contrast was set to 0.1 in the 1 c/d condition and 0.2 in the 5 c/d condition. Stimuli were Gabor patches of 5° size, 1 or 5 c/d spatial frequency, 45° or 135° orientation. In each trial, the stimulus was displayed for 500 ms in an half-period sinusoidal temporal envelope, stimulus presentation was accompanied by an auditory cue. The subject’s task was to identify the orientation of the Gabor patch: 45° or 135° by pressing keyboard keys. After response, audio feedback was provided. There was no fixation point on screen. Threshold were subsequently estimated by fitting a Weibull function to the data with a maximum likelihood procedure. The thresholds difference from baseline are reported in dB such that *Δtthresh*_*t*_ = 20*log_10_(*thresh*_*t*_/*thresh*_*0*_) where *thresh*_*t*_ is the measured threshold at given timepoint *t* and *thresh*_*0*_ is the measured threshold at baseline.

## Results

Contrast thresholds were measured over a period of one hour in a binocular control condition and in a monocular deprivation condition in 15 adults (Figure1a, see supplementary methods). Two spatial frequencies were tested 1 c/d and 5 c/d for a total of 4 experimental sessions, tested on different days in a pseudo-randomized counterbalanced order. In the monocular conditions, participants wore a translucent eye-patch on one of their eyes during the whole one-hour course of the experiment.

**Figure1:**
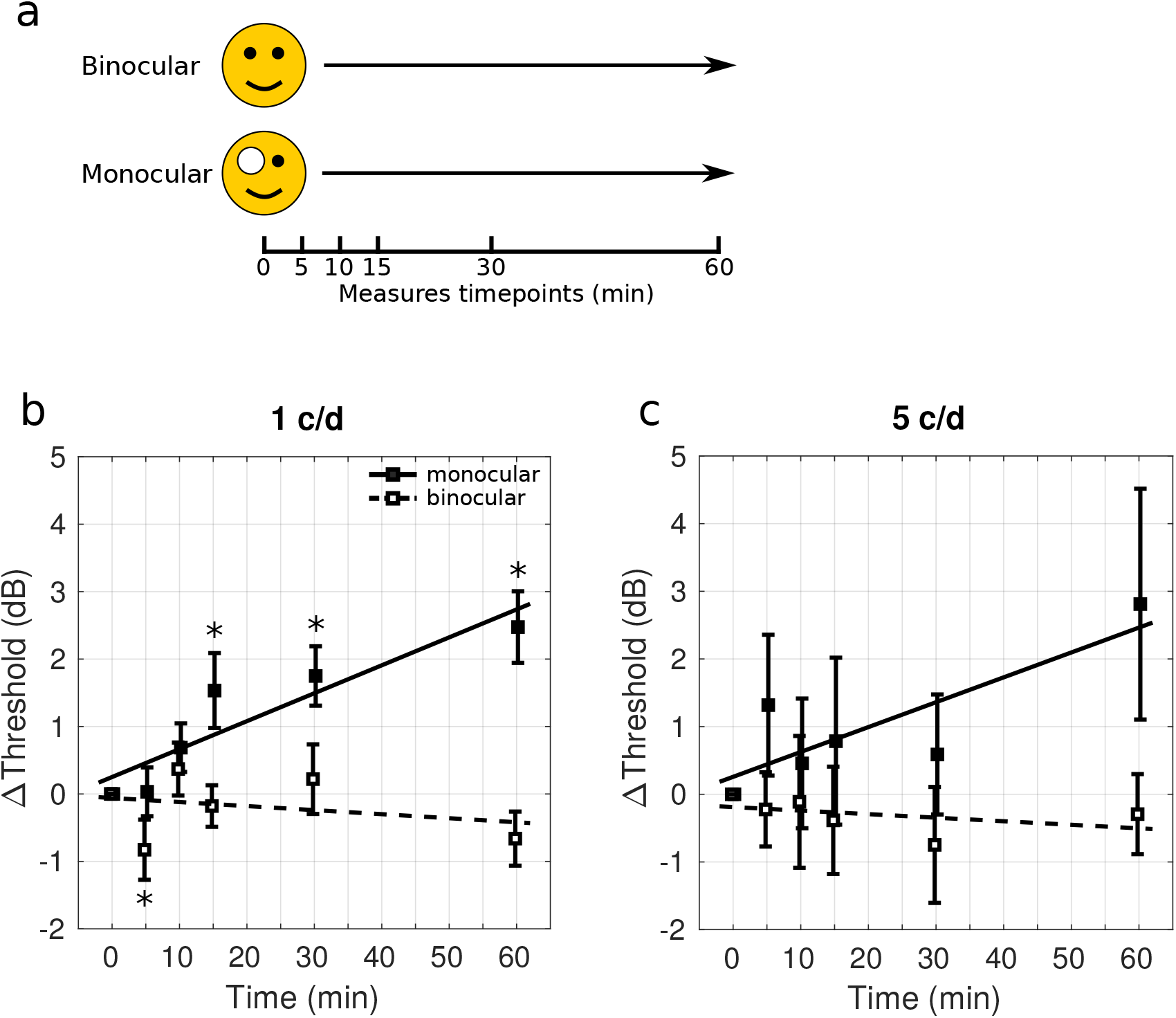
a) Protocol. Two conditions were tested: binocular i.e. the control condition or monocular i.e. the deprived condition. Threshold measurements were repeated at 0, 5, 10, 15, 30 and 60 minutes in the time course of the experiment. In the monocular condition, participants wore a translucent eye patch on one of their eyes during the whole one-hour course of the experiment. b) Results for a 1 c/d stimulus. Average threshold differences from initial baseline 0 min measurement at each measured timepoint for the binocular (open symbols) and monocular (closed symbols) condition. Errorbars indicate ± standard error, asterisks indicate significant difference from 0 dB (two-sided Wilcoxon signed-rank test, α=0.05). Regression lines of the average timepoints are plotted (continuous for the monocular and dashed for the binocular conditions). c) Same as b) tested with a 5 c/d stimulus.

As expected, in the binocular condition, for both 1c/d and 5c/d the contrast thresholds remain stable over time (Figure1b&c, open symbols and dotted lines). However, in the monocular deprivation condition at 1c/d, they increase over time (Figure1b, plain symbols and lines) and become significantly different both from baseline and from the binocular condition after 15 minutes of monocular deprivation (two-sided Wilcoxon signed-rank test, α=0.05). This monotonic increase seems to progressively slow-down, approaching a plateau after 30 minutes of deprivation. The increase in contrast threshold is also captured by the fact that both the Area Under the Curve (AUC) described by the thresholds time course and the linear regression of these thresholds over time are significantly positive in the monocular condition (one-sided Wilcoxon signed-rank test, p<0.001, the slope increases for all 15/15 subjects and the AUC is positive for 13/15 subjects, see Supplementary Figure1a&c) whereas they are not in the binocular condition. Furthermore, a two-way repeated measures ANOVA also reveals a significant interaction between the condition and time factors (p=0.001) indicating different time-courses in the two conditions.

For the 5 c/d condition, the trends are similar however less significant (Figure1c). In the monocular deprived condition, the contrast thresholds increase over time but they are not significantly different from either the baseline or from the binocular condition (two-sided Wilcoxon signed-rank test, α=0.05). Both the AUC described by the thresholds and the linear regression of the thresholds over time are not significantly different from 0 in either the monocular or binocular condition (one-sided Wilcoxon signed-rank test, α=0.05). However, the linear regressions are almost significantly positive in the monocular condition (p=0.084) and become significant when excluding one subject who shows a strong opposite trend because of a high baseline (p=0.045, see Supplementary Figure1b). The two-way repeated measures ANOVA doesn’t reveal any significant interaction between the condition and time factors (p=0.234). However, the time factor is significant, indicating a change in the threshold over time (p=0.006).

## Discussion

Zhou et al. (2013) reported an increase in the contrast threshold of the non-deprived eye, measured only after the deprivation period. Our results show that while one eye is patched, the contrast detection threshold of the other eye gradually increases during the period of deprivation particularly at lower spatial frequencies. This suggests that the contrast gain changes that result in a reciprocal change in the binocular balance (Zhou et al., 2013; Chadnova et al., 2017) build up slowly during the deprivation period as a result of the disrupted binocular input.

In the classical binocular combination models (Meese et al., 2006; Moradi and Heeger, 2009) the increase of monocular threshold without input in the other eye can be explained by one of three possible mechanisms: a reduction of the gain of the deprived eye, an increase in the normalization constant or a decrease in the slope of the psychometric function. The latter would represent an increased amount of noise which could be thought of as an intermittent interference/suppression from the patched eye. We ruled out this explanation by performing a control experiment in which we measured the full psychometric function with a constant stimuli procedure at the beginning and end of a one-hour patching period and showed that, although the threshold increased, the slope of the psychometric function for the non-patched eye did not become shallower over time (Supplementary Figure2). So far, electrophysiological studies (Chadnova et al., 2017; Binda et al., 2018) have observed a decrease in the neuronal population activity generated from the stimulation of the non-patched eye, after patch removal. Among the two remaining hypotheses, such a decrease would be inconsistent with a normalization explanation which would require increased neural population activity in the normalization pool (Moradi and Heeger, 2009; Busse et al., 2009; Reynaud et al., 2012). It is, however, consistent with a contrast gain reduction.

This study provides further insights into the plasticity phenomena involved in short-term monocular deprivation. These phenomena would be the consequence of contrast-gain changes that build up progressively during the period of deprivation. It also highlights a hitherto unknown feature of human vision, namely that monocular vision *per se* is intrinsically unstable which is a consequence of the reciprocal inhibitory circuits that homeostatically regulate binocular vision (Meese et al., 2006). This questions a whole corpus of studies of visual function that rely on the assumption that monocular vision is intrinsically stable.

