## Supplementary figures and images for "Monocular vision is intrinsically unstable: a side-effect of binocular homeostasis"

### FigureS1

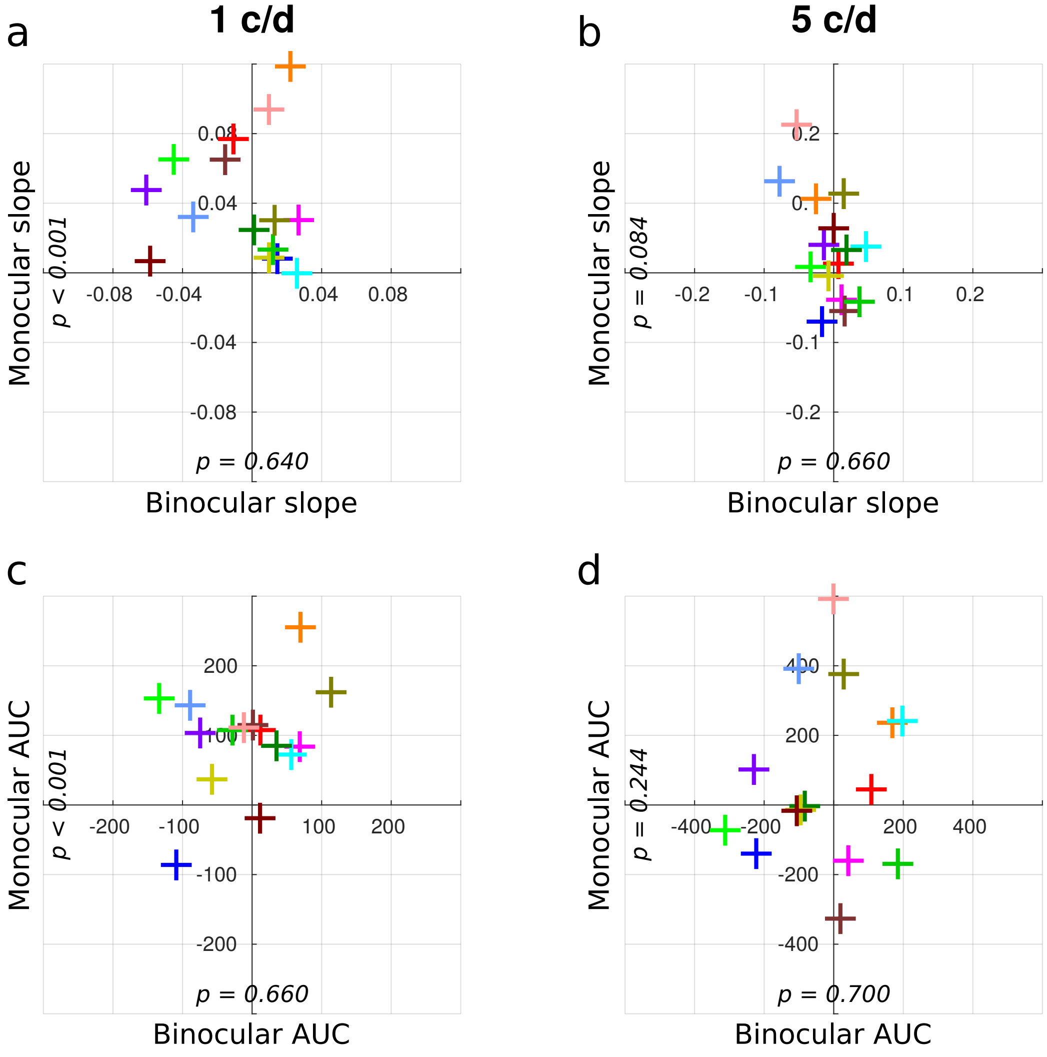


#### Supplementary Figure 1

### FigureS2

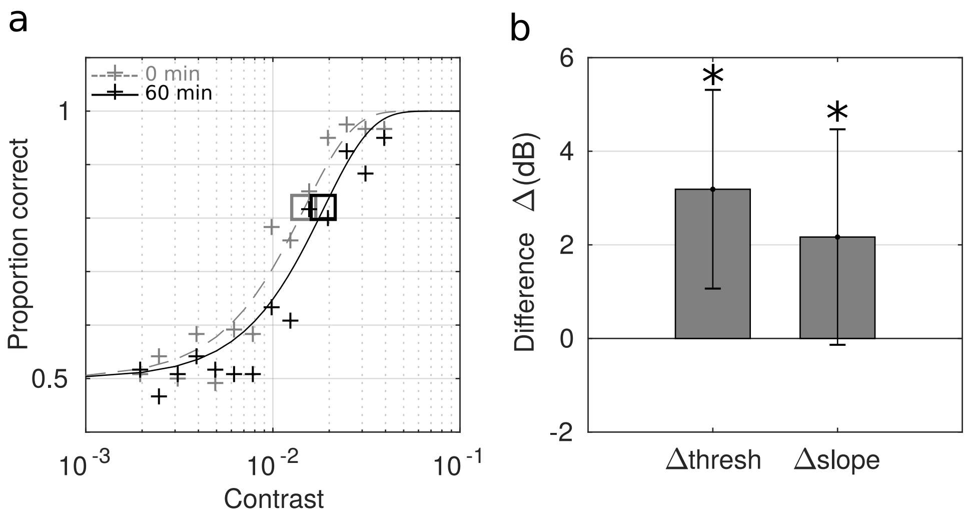


#### Supplementary Figure 2
