## SupplementaryCaptions for "Monocular vision is intrinsically unstable: a side-effect of binocular homeostasis"

**Supplementary materials**

Supplementary Figure 1: Scatterplots of the regression slopes (slopes in dB/min) of the evolution of the thresholds and their area under the curves (AUC in dB*min) for all the subjects in the monocular (y-axis) *vs*. binocular (x-axis) condition. a) regression slopes at 1 c/d. b) regression slopes at 5 c/d. c) AUC at 1 c/d. d) AUC at 5 c/d. Each symbol represents one individual subject (same color in all panels). p-values indicate significance levels of the one-sided Wilcoxon signed-rank test. The p-value of the slopes in the monocular condition at 5 c/d becomes p=0.045 when the subject identified by a brown symbol is excluded because of a high baseline.

Supplementary Figure 2: Control experiment. a) average psychometric functions of the detection of a 1 c/d Gabor patch measured with the method of constant stimuli for a subset of 7 participants at the beginning (0 min, gray dashed-line) and at the end of a 60 minutes (60 min, black continuous line) monocular deprivation period. Squares indicate thresholds at 82%. b) Difference in dB in the thresholds and slopes of the psychometric functions between the beginning and the end of the deprivation period. $\Delta thresh = 20*\log_{10}(thresh_{60}/thresh_{0})$ and $\Delta slope = 20*\log_{10}(slope_{60}/slope_{0})$ where $thresh_0$ and $thresh_{60}$ are the thresholds respectively measured at the beginning and the end of the deprivation period and $slope_0$ and $slope_{60}$ are the slopes of the psychometric function respectively measured at the beginning and the end of the deprivation period. Average between 7 subjects. Error bars represent standard deviation. Asterisks indicates values are significantly different from 0 dB (two-sided Wilcoxon signed-rank test, $\alpha$=0.05). Both the threshold and slope increased at the end of the monocular deprivation period. This indicates that the slope of the psychometric function for the non-patched eye did not become shallower over time, but actually became steeper.
